# Genome-wide Association Study of Anxiety and Stress-related Disorders in the iPSYCH Cohort

**DOI:** 10.1101/263855

**Authors:** Sandra M. Meier, Kalevi Trontti, Thomas Damm Als, Mikaela Laine, Marianne Giørtz Pedersen, Jonas Bybjerg-Grauholm, Marie Bækved-Hansen, Ewa Sokolowska, Preben B. Mortensen, David M. Hougaard, Thomas Werge, Merete Nordentoft, Anders D. Børglum, Iiris Hovatta, Manuel Mattheisen, Ole Mors

## Abstract

Anxiety and stress-related disorders (ASRD) are among the most common mental disorders with the majority of patients suffering from additional disorders. Family and twin studies indicate that genetic and environmental factors are underlying their etiology. As ASRD are likely to configure various expressions of abnormalities in the basic stress-response system, we conducted a genome-wide association study including 12,655 cases with various anxiety and stress-related diagnoses and 19,225 controls. Standard association analyses were performed supplemented by a framework of sensitivity analyses. Variants in *PDE4B* showed consistent association with ASRD across a wide range of our analyses. In mice models, alternations in *PDE4B* expression were observed in those mice displaying anxious behavior after exposure to chronic stress. We also showed that 28% of the variance in ASRD was accounted for by common variants and that the genetic signature of ASRD overlapped with psychiatric traits, educational outcomes, obesity-related phenotypes, smoking, and reproductive success.

## Introduction

Anxiety and stress-related disorders (ASRD) are among the most common mental disorders with a lifetime prevalence of over 20%^1^. In Europe alone, over 70 million people are annually suffering from these disorders^2^. Over 90% of people diagnosed with ASRD have another comorbid mental or somatic disorder^3^. Given the prevalence and the immense social and economic burden of ASRD^2^, it is of strong interest to identify their risk factors.

Family and twin studies indicate that both genetic and environmental factors are of relevance, with levels of familial aggregation and heritability at 30-50%^4^. As for other complex genetic disorders, many linkage and candidate gene association studies have been conducted, but with little success in the robust identification of susceptibility genes for ASRD. Genome-wide association studies (GWAS) have proven to be an effective tool for the identification of common genetic variants increasing the susceptibility to complex disorders. Recently, GWAS of ASRD such as panic disorder^5^ߝ^8^, post-traumatic stress disorder (PTSD)^9–15^, generalized anxiety disorders^16^, phobias^17, 18^, and a composite indicator of anxiety disorders^19, 20^ have been published. However, these efforts have been limited by small sample sizes resulting in low overall power to detect significant associations.

ASRD are likely to configure various expressions of abnormalities in the basic stress-response system. In line with this hypothesis, twin studies found a strongly shared genetic susceptibility^21, 22^ Accordingly we aimed to conduct a GWAS aggregating cases in the Lundbeck Foundation Initiative for Integrative Psychiatric Research (iPSYCH) study with varying diagnoses of ASRD. As most people with ASRD are suffering from another comorbid mental disorder, especially depression^23^; we explored the impact of mental comorbidity on the genetics of ASRD. Our effort represents the first genetic study of this magnitude to explicitly target comorbidity of ASRD with other mental disorders.

## Results

### GWAS and gene-based analysis

In total, 68 genetic variants in a single locus exceeded the threshold for genome-wide significance in our “naïve” GWAS for ASRD, i.e. without correction for the study design. The locus overlaps with one gene (Phosphodiesterase 4B (*PDE4B*)) with the top SNP being rs7528604 (P=5.39*10^−11^, OR=0.89, Figure 1, Table 1, Figure S1). We found no evidence for significant heterogeneity between genotyping batches for this marker (Figure S2). Stratified analyses focusing on anxiety and stress-related disorder phenotype definitions separately supported the identified locus (Figure S3 and S4). In addition, results from analyses mimicking the comorbidity pattern of population-based samples (rs7528604, P=1.20*10^−11^, OR=0.88) and including psychiatric phenotypes as covariates (rs7528604, P=2.32*10^−8^, OR=0.90) were in line with our initial GWAS results (Figure S5 and S6). Even in the stringent sensitivity analyses, associations within the *PDE4B* gene were among the top signals (rs17128482, P=1.43*10^−5^, OR=1.12, Figure S7). Resampling analyses removing comorbid depression cases from our data did not show a significant impact for depression on our result at the *PDE4B* locus (P=0.464). Downstream analyses using both GTEx^24^ and the SNP tag lookup function in MR-Base (www.mrbase.org/beta) indicated that there is currently no evidence for the top SNPs to be, directly or via LD tagging, expression or methylation quantitative trait loci.

**Figure 1.**
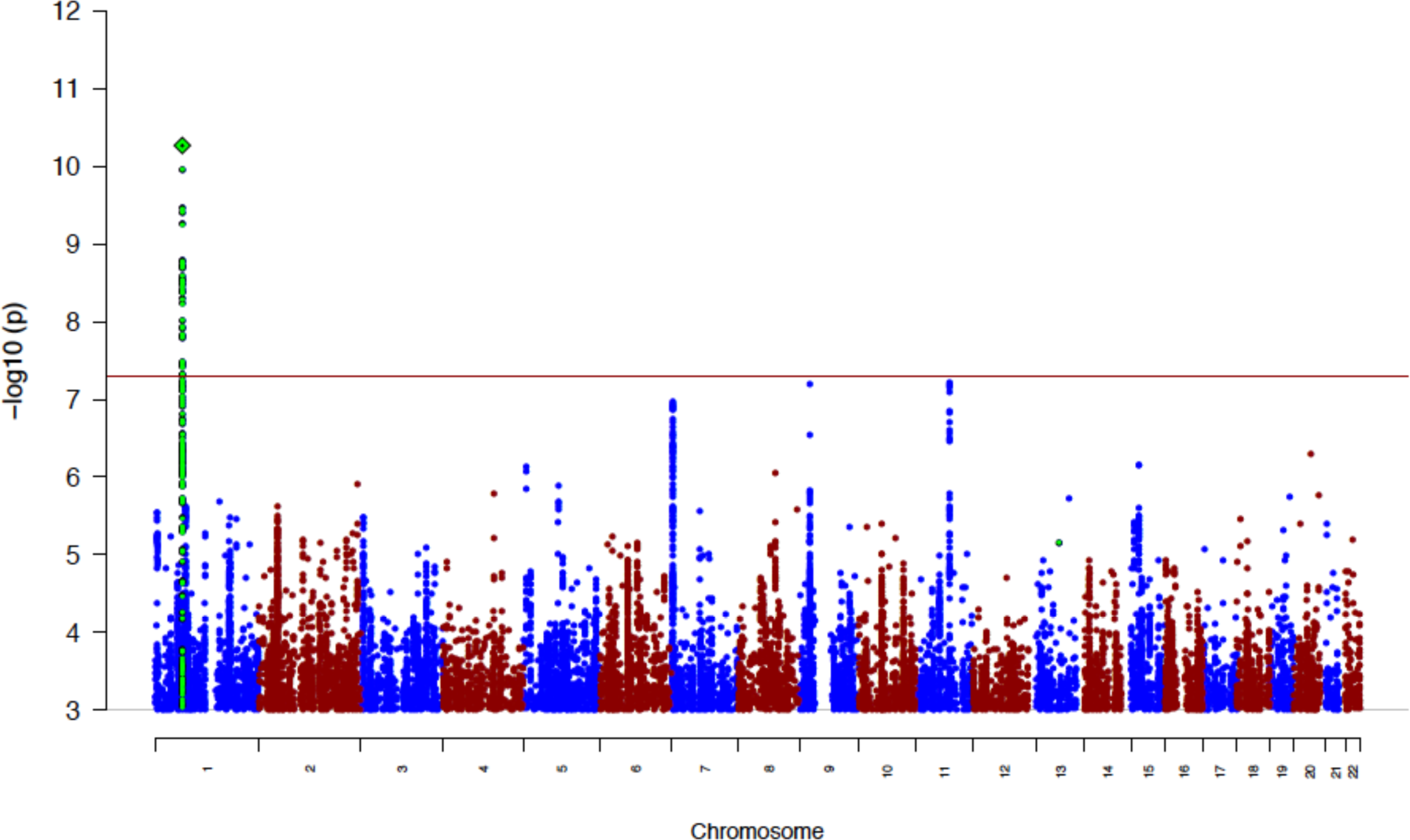
Manhattan plot of the results from the GWAS of anxiety and stress-related disorders. The index variants in the most significant loci are highlighted as a diamond. Index variants located with a distance less than 400kb are considered as one locus.

**Table 1.** Results for index variants in the top 10 loci associated with anxiety and stress-related disorders. Index variants are LD independent (r^2^ < 0.1), and are merged into one locus when located with a distance less than 400kb. The location (chromosome [Chr] and base position [BP]), single nucleotide polymorphism (SNP), alleles (A1 and A2), allele frequency (A1 Freq), odds ratio (OR) of the effect with respect to A1, and association P-value of the index variant are given, along with genes within 100kb of the locus.

| Locus | CHR | BP | SNP | Genes | A1 | A2 | A1 Freq | OR | P-value |
| --- | --- | --- | --- | --- | --- | --- | --- | --- | --- |
| 1 | 1 | 66407352 | rs7528604 | PDE4B | A | G | 0.385 | 0.890 | $5.39 \times 10^{-11}$ |
| 2 | 11 | 81047274 | rs1458103 | - | A | C | 0.741 | 0.898 | $6.19 \times 10^{-8}$ |
| 3 | 9 | 2511193 | rs113209956 | - | T | C | 0.085 | 0.828 | $6.36 \times 10^{-8}$ |
| 4 | 7 | 3676002 | rs6462203 | SDK1 | A | C | 0.265 | 0.901 | $1.09 \times 10^{-7}$ |
| 5 | 20 | 41070559 | rs6030245 | PTPRT | T | C | 0.795 | 1.120 | $5.06 \times 10^{-7}$ |
| 6 | 15 | 41024303 | rs11855560 | CASC5, LOC100505648, RAD51, RMDN3, GCHFR, DNAJC17, C15orf62, ZFYVE19 | T | C | 0.469 | 1.089 | $6.96 \times 10^{-7}$ |
| 7 | 5 | 7448796 | rs2451828 | ADCY2 | T | C | 0.019 | 1.340 | $7.37 \times 10^{-7}$ |
| 8 | 8 | 87643741 | rs16916239 | CNGB3 | A | G | 0.783 | 0.903 | $8.96 \times 10^{-7}$ |
| 9 | 2 | 233649290 | rs79928194 | GIGYF2, KCNJ13 | T | C | 0.905 | 0.862 | $1.26 \times 10^{-6}$ |
| 10 | 5 | 83470986 | rs342422 | EDIL3 | A | G | 0.53 | 0.920 | $1.28 \times 10^{-6}$ |

Gene-wide analyses using MAGMA^25^ identified seven genome-wide significant genes, six of which did not co-localize with our genome-wide significant SNP locus at *PDE4B* (P_gene_=4.74*10^−15^): *SDK1* (P_gene_=5.79*10^−7^) on chromosome 7, and *GCHFR* (P_gene_=3.30*10^−7^), *RMDN3* (P_gene_=9.36*10^−7^), *C15orf62* (P_gene_=1.76*10^−6^) and *RAD51* (P_gene_=2.58*10^−6^), on chromosome 15 sharing their top SNPs. No pathways were significant after correction for multiple testing (Table S3 and S4). Besides *PDE4B* no previously described candidate gene reached gene-based significance (for a detailed review of the candidate gene literature see elsewhere ^26, 27^).

### SNP heritability and genetic correlation with other traits

LD score regression^28, 29^ was used to calculate SNP heritability of our “naïve” ASRD GWAS and the genetic correlation with other phenotypes. Assuming a population prevalence of 20% for ASRD, we estimate that the liability-scale SNP heritability was 0.28 (SE=0.027). These estimates are comparable to those reported in previous studies for PTSD^30^, but higher than for anxiety disorders^19^. Partitioning heritability based on functional annotations revealed significant enrichment in the heritability by SNPs located in conserved regions (enrichment=2.01, SE=0.32, P=0.0025), supporting the general biological importance of conserved regions and their potential impact on susceptibility of ASRD (Figure 2). Cell-type specific analyses revealed a significant enrichment in the heritability by SNPs located in central nervous system specific enhancers and promoters (enrichment=2.93, SE=0.50, P=4.32*10^−4^, Figure S8 and S9).

**Figure 2.**
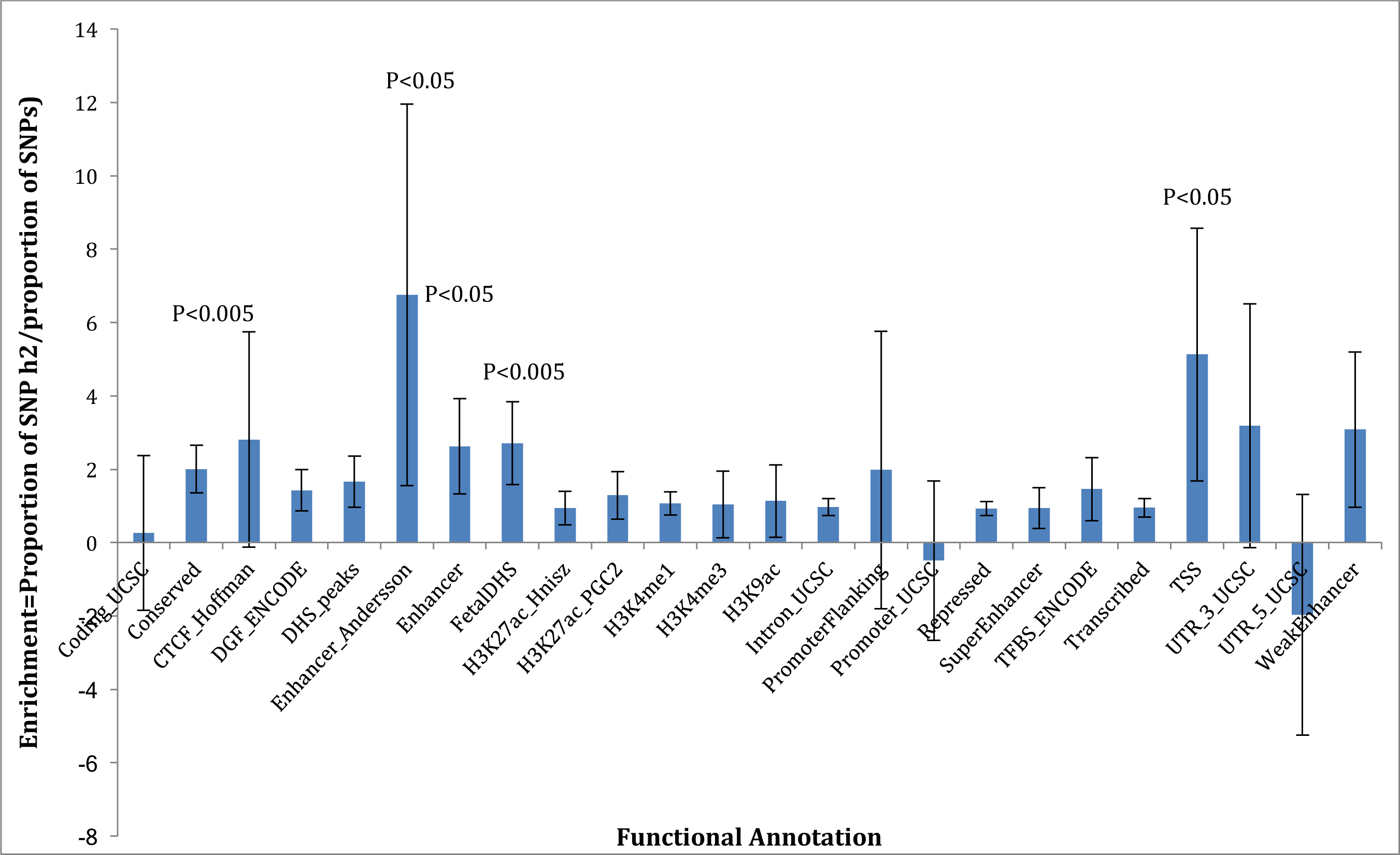
Partitioning of heritability by functional annotations. Enrichment of heritability per SNP in 24 functional annotations defined by Finucane et al.^38^ Error bars represent 95% confidence intervals. P-values for annotation categories with nominal significant enrichment are shown.

Using LD Hub^31^ 31 phenotypes displayed significant genetic correlation with ASRD after Bonferroni correction (P=2.19*10^−4^), including psychiatric traits, educational outcomes, obesity-related phenotypes, smoking, and reproductive success (see Figure 3). Notable significant genetic correlations between anxiety and stress-related and psychiatric traits were identified for depressive symptoms (r_g_=0.73, P=6.99*10^−20^), subjective well-being (r_g_=−0.46, P=3.53*10^−10^), major depressive disorder (r_g_=0.58, P=7.20*10^−9^), neuroticism (r_g_=0.49, P=1.65*10^−7^), and schizophrenia (r_g_=0.30, P=1.57*10^−12^). Positive genetic correlations were also found for traits related to obesity including significant relationships with body mass index (r_g_=0.17, P=8.85*10^−6^), waist-to-hip ratio (r_g_=0.19, P=1.05*10^−5^), and body fat (r_g_=0.23, P=8.72*10^−5^). An overview of significant genetic correlations for ASRD can be found in Tables S5 and S6. LD score regression (LDSC)^28, 29^ revealed a strong genetic correlation (r_g_=0.46, P=0.0021) with the ANGST Consortium^19^. We performed the same LDSC-based analyses also for the results from our framework of sensitivity analyses. The impact of the analyses mimicking the comorbidity pattern of population-based samples on SNP heritability and the genetic correlations with other traits was marginal (see Tables S7 and S8). Due to the overrepresentation of psychiatric phenotypes in the control sample the design including psychiatric phenotypes as covariates and the propensity score matched design resulted in lower SNP heritability and genetic correlations (see Tables S9-12). The interpretability of these findings is therefore not straightforward, however, results from these analyses might provide further insight into the nature of the comorbidity of ASRD with other psychiatric phenotypes. An overview with regard to SNP heritability reflecting the increased prevalence of ASRD in psychiatric phenotypes can be found in Table S13.

**Figure 3.**
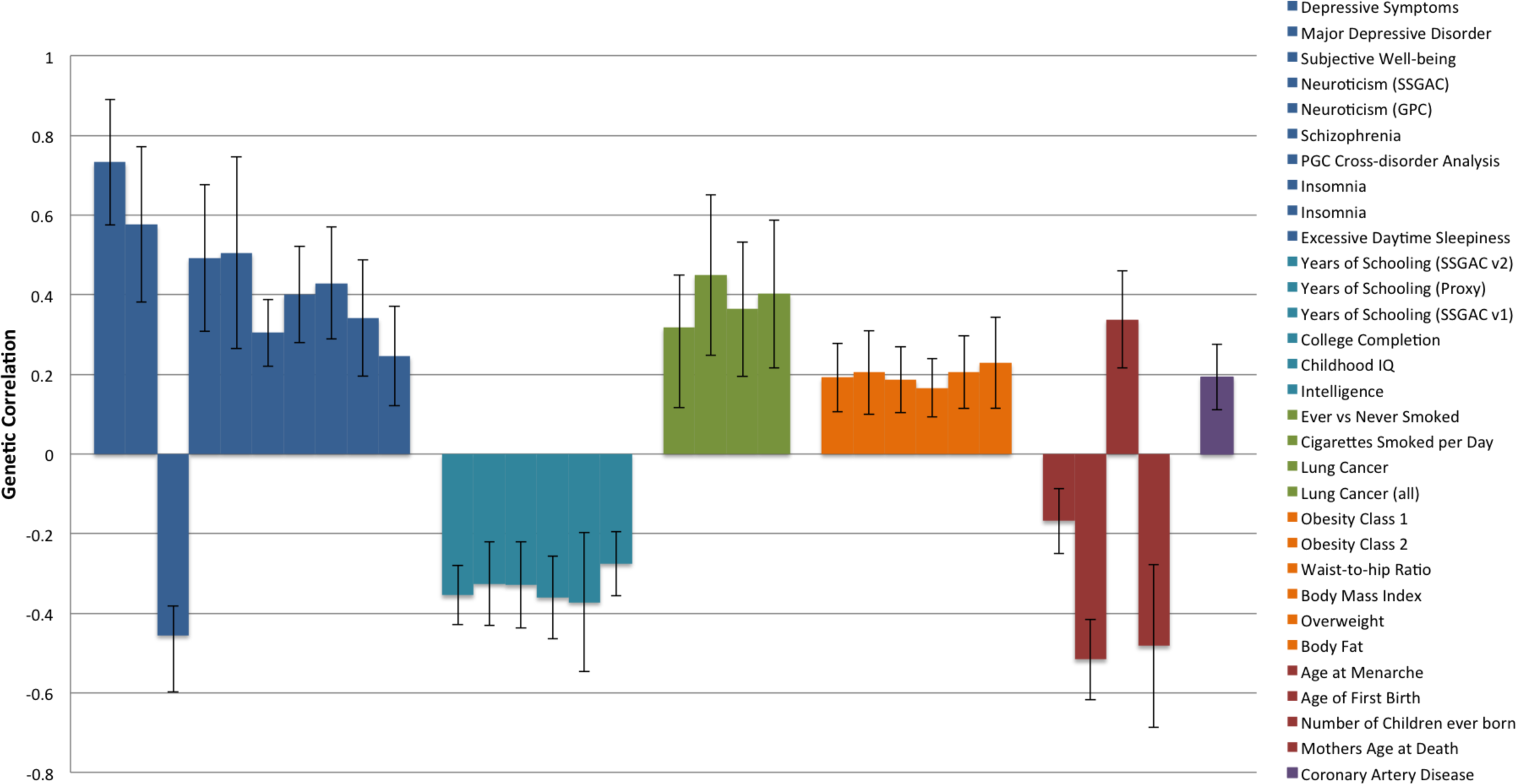
Significant genetic correlations between anxiety and stress-related disorders and other heritable traits. In total 228 traits were tested (including psychiatric traits, educational outcomes, obesity-related phenotypes, smoking, and reproductive success). Error bars indicate 95% confidence limits.

### Polygenic risk score

PRS computed in our study using the case-control GWAS results from the ANGST Consortium^19^ as the discovery sample were in a dose-response relationship associated with higher OR for ASRD (Figure S10). The maximum variance explained by the estimated PRS (Nagelkerke’s R^2^) was 0.2% (SE=5.36*10^−4^) at P-value threshold of 0.1, which is in line with observations in similarly sized GWAS for depression and anxiety disorders. The odds ratio in standardized PRS (10^th^ decile) between cases and controls was 1.33 (95%CI: 1.20-1.48).

### Our results in context

In this section we compare our GWAS with previous work related to these phenotypes^19, 20, 30, 32^. Using MAGMA^25^, we replicated associations within the *CAMKMT* gene (P_gene_=2.97*10^−4^) identified by the ANGST Consortium^19^, with the lead SNP displaying a different direction of effect (rs1067327, P=1.59*10^−4^, OR=0.93). There were no genome-wide significant SNPs or genes reported in the PTSD meta-analysis^48^. SNPs identified in an UKBB study were significantly associated with anxiety (rs3807866, P=0.048, OR=1.05) and stress-related disorders separately (rs10809485, P=0.042, OR=1.04). With regards to our own finding for *PDE4B* the results for the ASRD spectrum in other samples are diverse. *PDE4B* was not significantly associated with anxiety disorders in the ANGST Consortium (P_gene_=0.554, lead SNP rs7528604, P=0.884, beta=0.0039). However, *PDE4B* was associated with PTSD, although with a different direction of effect (rs7528604, P=0.010, OR=1.10). Initial genome-wide screenings within UK biobank for nerves, anxiety, tension and depression seem to support *PDE4B* as susceptibility locus (rs7528604, P=0.0057, beta=−0.0032, https://sites.google.com/broadinstitute.org/ukbbgwasresults/). Furthermore, our lead SNP rs7528604 has been found to be associated with neuroticism (P=0.0011, beta=−0.0181)^32^, which is often used as a proxy for ASRD.

Recently the largest GWAS for schizophrenia to date provided evidence for the association of *PDE4B* with schizophrenia at the level of genome-wide significance^33^. It is therefore of no surprise that our lead SNP rs7528604 is associated with schizophrenia (P=0.0023, beta=−0.033)^34^. In order to assess, whether our *PDE4B* signal might have been triggered by cases with comorbid schizophrenia in the iPSYCH cohort, we excluded cases with schizophrenia and still observed genome-wide significance for our lead SNP (P=3.58*10^−10^, OR=0.89).

### Lower *Pde4b* expression levels in mice

We determined the gene expression levels of *Pde4b* in mice exposed to CSDS using RNA-Seq (Figure 4). Stress susceptible mice from the B6 strain had lower expression levels of *Pde4b* in the mPFC compared to both control (P=0.002, P_adj_=0.063) and stress resilient mice (P=0.006, P_adj_=0.098), as well as in the vHPC compared to control (P=0.001, P_adj_=0.004) and stress resilient mice (P=0.003, P_adj_=0.014). D2 mice are highly susceptible to CSDS and therefore, we were only able to compare stress susceptible mice to controls. There were no differences in *Pde4b* expression levels between these groups in either brain region.

**Figure 4.**
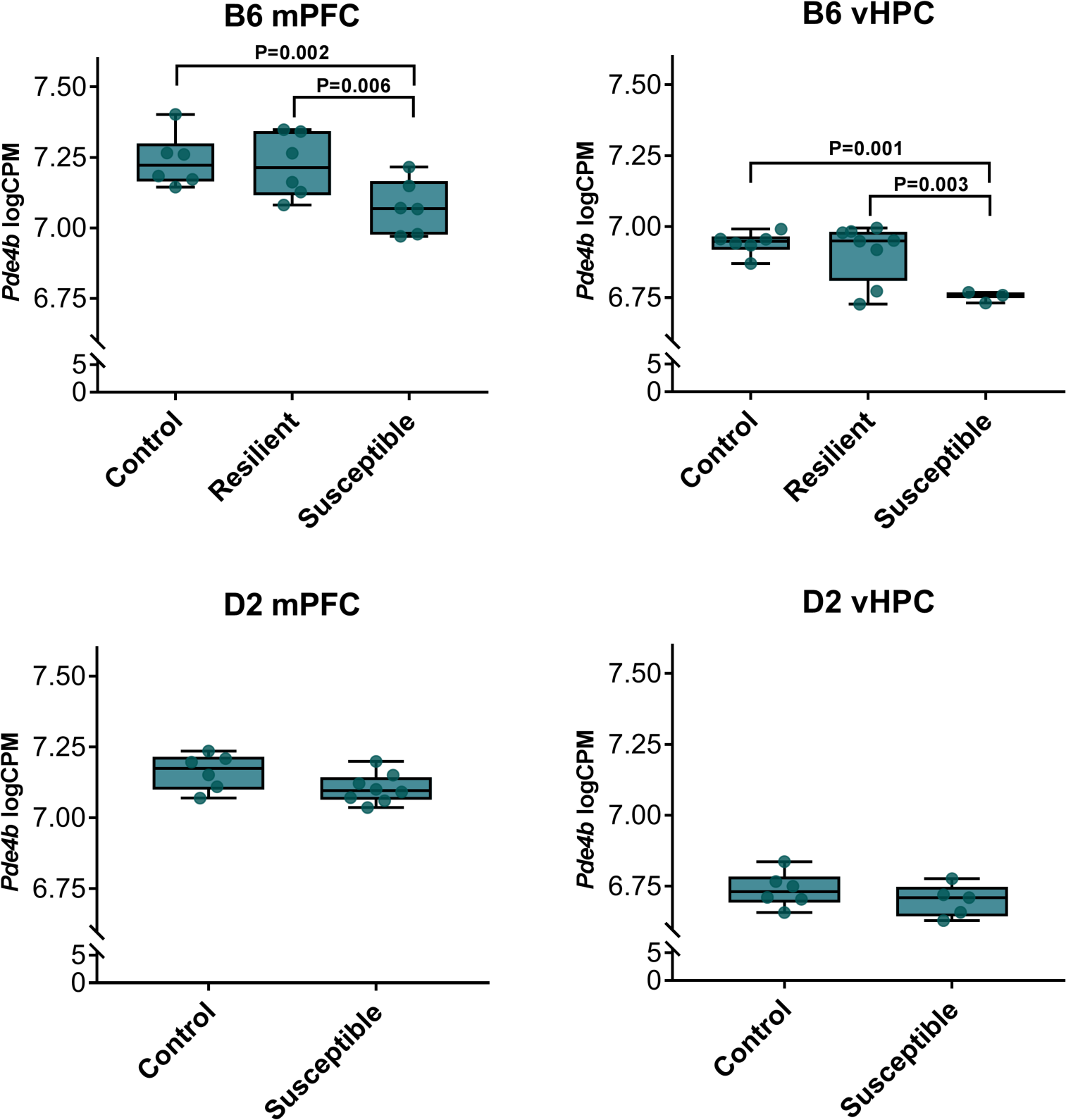
Differential *Pde4b* gene expression levels in mice exposed to chronic psychosocial stress. Each sample is represented by a dot superimposed on the box plot. Number of mice / group: B6 strain Controls = 6 (mPFC & vHPC), Resilient = 6 (mpFC) and 8 (vHPC), Susceptible = 6 (mpFC) and 3 (vHPC), D2-strain Controls = 6 (mpFC & vHPC), Resilient = 0, Susceptible = 8 (mpFC) and 5 (vHPC).

## Discussion

To the best of our knowledge, we conducted the largest GWAS on ASRD to date and extend previous findings about shared genetic effects with other psychiatric and somatic phenotypes. Specifically, we aggregated 12,655 cases with varying clinical diagnoses of ASRD and 19,225 control subjects with the aim of identifying common variants underlying the etiology of these disorders. While there is room for improvement, we tried to capture the clinical complexity of anxiety and stress-related phenotypes in a sample enriched for individuals with comorbid mental illnesses and identified genetic variants that were associated with disease susceptibility.

The most consistent association signal across our different analyses (albeit not genome-wide significant in the propensity score matched design) was observed for genetic variants located within the *PDE4B* gene, which regulates intracellular cAMP signaling and is strongly expressed in the human brain. *PDE4B* has been proposed as a candidate gene for anxiety, in particular panic disorder^35^, as mice deficient in *Pde4b* exhibit behavioral changes in range of tests sensitive to anxiolytic drugs^36^. Consistent with the behavioral data, plasma corticosterone concentrations were increased in these mice^36^. We found the expression of *Pde4b* to be altered in B6 mice susceptible to chronic psychosocial stress compared to controls and stress-resilient mice. Lower expression levels were observed in both examined brain regions (mPFC and vHPC), which are known to regulate emotional and social behavior in mice and humans. In innately anxious (D2 inbred) mice, *Pde4b* expression did not differ from controls after the chronic psychosocial stress exposure indicating a genetic background effect. This finding is in line with anxiety being a complex phenotype involving the interplay of environmental stressors and multiple genes. Additionally, pharmacological profiles of selective *PDE4B* inhibitors have demonstrated clear antidepressant and anxiolytic benefits^37^.

In line with other GWAS^19, 30^, our SNP heritability estimate of 28% for ASRD indicates a substantial role for common genetic variation, which accounts for a sizable portion of twin-based heritability^4^. Evolutionarily conserved regions and regions containing enhancers and promoters of expression in the central nervous system tissues were found to be enriched for associations with ASRD, consistent with findings for schizophrenia, bipolar disorder, and depression^38^. More work is needed to unravel the nature of the genetic correlations described in our manuscript and how different analysis designs in our analytical framework impacted these findings. Nevertheless, the range of genetic correlations with psychiatric traits, educational outcomes, obesity-related phenotypes, smoking, and reproductive success is in itself remarkable and helps to broaden our conceptualization of ASRD. First, the strong positive genetic correlations of ASRD with depression and neuroticism in our “naïve” GWAS reinforce clinical and epidemiological observations. ASRD are commonly comorbid with depression, often precede depression and even affect the course of the depression. Second, the positive genetic correlations seen with schizophrenia and the cross-psychiatric disorder phenotype firmly anchor ASRD with other psychiatric traits and reflect the substantial evidence for partially shared genetic susceptibility across many psychiatric disorders^39^. Third, negative associations between ASRD and educational attainment have been reported^40^, and our results suggest that genetic factors may partially account for these reported associations. Fourth, the identification of significant positive correlations with obesity-related phenotypes are in concordance with unusual high prevalence of anxiety disorders in morbidly obese patients.^41^

A major strength of this study is the aim to identify genetic variants that play a central but nonspecific role in the susceptibility of ASRD. This contrasts with the approach taken in most psychiatric genetic studies, which generally focus on specific clinical diagnoses. However, it has long been recognized that clinical diagnoses have their shortcomings and poorly reflect etiological mechanisms, as both genetic and environmental factors have been found to have non-specific effects across a wide range of diagnoses, particularly among anxiety disorders^21^. Given how critical fear and anxiety are for human survival, it is very likely that highly conserved genes common to a range of anxiety and stress-related disorder regulate these basic biological processes. Making use of registered-based diagnoses as a proxy for mental disorders in a research study (iPSYCH) that was primarily ascertained for closely related traits (such as depression and others) constitute the major limitation, but also strength of our study enabling the characterization of ASRD susceptibility in the context of mental comorbidity. Similar to previous studies in ASRD the generalizability to milder forms of ASRD or truly population-based samples is difficult to assess. Through different sensitivity analyses we aimed to address some of the limitations (to the extent possible with the data at hand) and gained new insights that probably would not have been possible otherwise. It is of note, that despite this lack in generalizability the above-mentioned limitations are unlikely to lead to false positive associations in a narrow sense. They do however ask for a reflection on the tested hypothesis in the (sensitivity) analyses under consideration. As a final point we would like to stress that our replication efforts were limited by the fact that our study is including a wider range of both ASRD than previous efforts^8, 20, 30^, which were still restricted in their sample size and power.

In summary, our results highlight ASRD to be a complex heritable phenotype with intriguingly large and significant genetic correlations not only with psychiatric traits, but also with educational outcomes and multiple obesity-related phenotypes, thereby supporting established hypotheses about the biology of anxiety. Furthermore, we highlight the candidate gene *PDE4B* as a robust locus for ASRD (through studies in mice and men), pinpointing to the potential of *PDE4B* inhibitors in treatment of ASRD. Future studies are needed to confirm these findings via independent replication and to detect additional loci, not only identifying potential pleiotropic effects across the full spectrum of ASRD (as targeted in this study) but also loci associated specifically with each particular anxiety and stress-related disorder.

## Methods

### Sample

The GWAS sample under analysis included 12,655 cases with ASRD as well as 19,225 controls from Denmark and was identified as follows: All study participants were enrolled in iPSYCH, a study designed to unravel risk factors of severe mental disorders, including > 50,000 cases with schizophrenia, autism, attention-deficit/hyperactivity disorder (ADHD), anorexia nervosa, and affective disorders (referred to in this paper as “design” phenotypes) as wells as > 25,000 population-based controls. More information on iPSYCH can be found elsewhere^42^. 4,584 cases were diagnosed with an anxiety disorder, 9,831 with a stress-related disorder, of which 1,760 received both diagnoses. In brief, DNA samples for iPSYCH were taken from the Danish Neonatal Screening Biobank (DNSB)^43^. Following protocols for DNA extraction and amplification (described elsewhere^44^), all samples were genotyped using Illumina PsychChip (Illumina, San Diego, CA, USA). Through the national research registers we identified ASRD diagnoses assigned by psychiatrists according to the International Classification of Diseases 10th Revision (ICD10; (F40.0-F41.9; F43.0-F43.9); Table S1). Cases with an ASRD diagnosis and comorbid autism were excluded. Although patients with autism experience anxiety, their anxiety is often reflecting their core autistic symptomatology and lacks the social component central to many ASRD diagnoses^45^. Exclusion criteria for control subjects were ICD10 diagnoses of anxiety, stress-related as well as mood disorders. The Danish Data Protection Agency and the Danish Scientific Ethics Committee approved the study.

### Quality control, GWAS and gene-based analysis

Quality control, imputation and primary association analyses in iPSYCH have been described elsewhere^46^. In brief, we used the bioinformatics pipeline Ricopili (available at https://github.com/Nealelab/ricopili) developed by the Psychiatric Genomics Consortium (PGC)^34^. In order to avoid potential study effects of the 23 genotyping batches within the iPSYCH cohort, each batch was processed separately. Standard procedures for stringent quality control included filters for call rate, Hardy-Weinberg equilibrium, and heterozygosity rates. Each cohort was then phased and imputed using the 1000 Genomes Project phase 3 imputation reference panel^47^ using SHAPEIT^48^ and IMPUTE2^49^, respectively. Cryptic relatedness and population structure were assessed on high-quality single nucleotide polymorphisms (SNPs) with low linkage disequilibrium (LD) passing filters.

GWAS for the 23 genotyping batches in iPSYCH were performed using logistic regression models with the imputed marker dosages including the first four principal components in order to control for remaining population stratification^50^. Subsequently the results were meta-analysed using an inverse-variance weighted fixed effects model, implemented in METAL^51^. Additionally, the analyses were stratified for anxiety disorders and stress-related disorders. The majority of cases with ASRD were also diagnosed with another mental disorder (Table S2) and have been included in our study due to their comorbidity with one of the “design” phenotypes in iPSYCH. We therefore aimed to adjust for mental comorbidity in a framework of sensitivity analyses. With regards to the study design we ran two additional models, one that randomly selected a subset of ASRD cases mimicking the comorbidity patterns observed in population-based samples (following the idea of a “weighted design”) and a second that included psychiatric phenotypes as covariates^52^. We also removed individuals with comorbid depression in an attempt to identify the impact of this subgroup on the overall association. In our most stringent sensitivity analyses, we further employed a conservative propensity-score-matched design (see Supplemental Information).

Gene-based associations were calculated with MAGMA^25^ using the summary statistics from the main GWAS analyses. Association was tested using the SNP-wise mean model, in which the sum of −log (SNP P-value) for SNPs located within the transcribed region was used as test statistic. MAGMA^25^ controls for gene-size, number of SNPs in a gene, and LD between markers estimated from the 1000 Genomes Project phase 3.

### SNP heritability and genetic correlation with other traits

LD score regression was used to dissect the relative contribution of polygenic effects and confounding factors, such as cryptic relatedness, sample overlap and population stratification, to deviation from the null in the genome-wide distribution of GWAS χ2 statistics. Prevalence for calculation of liability scale heritability was specified as 20% for the combination of ASRD^53, 54^. Using LD score regression SNP heritability was also partitioned by functional category and tissue association^28, 29^. Partitioning was performed for 53 overlapping functional categories as well as 220 cell-type-specific annotations grouped into 10 cell-type categories^38^. Furthermore, genetic correlations (r_g_) were tested for 228 phenotypes with publically available GWAS summary statistics and 596 traits from the UK Biobank study (UKBB, https://sites.google.com/broadinstitute.org/ukbbgwasresults), using LD Hub^31^.

### Polygenic risk score

Polygenic risk scores (PRS)^55^ were constructed in our sample based on SNP-level summary statistics from the ANGST Consortium^19^ using variants passing a range of association P-value thresholds. PRS were calculated by multiplying the natural log of the odds ratio (OR) of each variant by the allele-dosage (imputation probability) and whole-genome PRS were obtained by summing values over variants for each individual^55^. For each target group and for each P-value threshold the proportion of variance explained (i.e. Nagelkerke’s R^2^) was estimated by comparing the regression with PRS to a reduced model with the same covariates only, but with the polygenic risk score term removed.

### Mouse model of chronic psychosocial stress

To establish the effect of chronic psychosocial stress on the brain gene expression levels of significant genes, we employed the chronic social defeat stress (CSDS) model^56^. Mice were exposed to a resident aggressor mouse (Clr-CD1 strain) for max. 10 min followed by housing in adjacent compartments for 24 h, repeated once per day for 10 days with a novel resident aggressor. Control mice were housed adjacent to another mouse, switching cage-mates daily but without physical exposure^56^.

Twenty-four hours after the last CSDS session we tested all mice for social aversion comparing their explorative behavior in the area around a cylinder with and without a Clr-CD1 mouse. The ratio of time the mouse spent in the area in the trial containing the mouse compared to the trial without a social target was used to determine whether each defeated mouse was resilient or susceptible to CSDS.

One week after the end of CSDS we dissected the medial prefrontal cortex (mPFC) and ventral hippocampus (vHPC) for RNA sequencing (RNA-Seq). All animal procedures were approved by the Regional State Administration Agency for Southern Finland (license numbers ESAVI-3801-041003-2011 and ESAVI/2766/04.10.07/2014) and carried according to directive 2010/63/EU of the European Parliament and of the Council, and the Finnish Act on the Protection of Animals Used for Science or Educational Purposes (497/2013).

### Gene expression profiling

Total RNA was extracted using TriReagent. For RNA-seq we carried out rRNA depletion using Ribo-Zero (Illumina, San Diego, CA, USA) followed by fragmentation with an S2 ultrasonicator (Covaris Inc., Woburn, MA, USA). mRNA sequencing libraries were prepared with Nextera (Illumina, San Diego, CA, vHPC samples) or ScriptSeq v2 (Epicentre, Madison, WI, USA, mPFC samples) kits. Following quality control, we aligned the mRNA reads with STAR 2.5.0c^57^ to the mouse genome GRCm38. Data normalization and differential expression analysis were carried out with ComBat^58^ and limma v3.32.5^59^. Whole transcriptome level multiple testing correction was done with the Benjamini-Hochberg method^60^, after which the expression levels of significant genes were extracted from the dataset.

## Acknowledgements

The iPSYCH team acknowledges funding from the Lundbeck Foundation (grant no R102-A9118 and R155-2014-1724), the Novo Nordisk Foundation for supporting the Danish National Biobank resource, and grants from Aarhus and Copenhagen Universities and University Hospitals, including support to the iSEQ Center, the GenomeDK HPC facility, and the CIRRAU Center. We thank Naomi Wray and Jack Hettema for helpful comments and suggestions during the preparation of the manuscript.

The Hovatta lab acknowledges funding from the European Research Council Starting Grant GenAnx, and thanks Ingrid Balcells, Natalia Kulesskaya, Suvi Saarnio, Jenni Lahtinen, and Sanna Kängsep from the Hovatta lab for technical help and discussions on mouse experiments. Mouse behavioral experiments were carried out at the Mouse Behavioural Phenotyping Facility, and RNA-seq at the The FIMM Technology Centre and The DNA Sequencing and Genomics Laboratory supported by University of Helsinki (HiLIFE) and Biocenter Finland.

## Author Contributions

S.M.M, M.M., and O.M. conceived the idea of the study. M.M. supervised the pre- and post GWAS analysis pipeline. S.M.M, T.D.A, and M.M. performed the analyses. K.T., M.L., E. S., and I. H. conducted experiments in mice. M.G.P, J.B.G, M.B.H., P.B.M., D.M.H, T.W., M.N., A.D.B., and O.M. provided and processed samples. S.M.M. and M.M. wrote the manuscript. All authors discussed the results, and approved the final version of the manuscript.

